# The Effect of Targeted Vaccination Against Mycobacterium Avium ssp. Paratuberculosis (MAP) in a Multiple Sclerosis Mouse Model: Implications for Causation

**DOI:** 10.1101/202275

**Authors:** Michael R. Warmoth

## Abstract

Epidemiologic evidence relating to the causation of Multiple Sclerosis, based upon twin concordance studies, implicates both genetic and environmental contributions. The HLA-DRB1/ HLA-A haplotype confers a 23 fold increase in MS prevalence above its baseline of 1 per 1,000 persons. Epigenetic factors, such as an aberrant response to Epstein Barr Virus (EBV) and Vitamin D deficiency can increase that risk 36 and 2 fold respectively. Evidence of an association between elevated MAP antibodies and Multiple Sclerosis has been reported.

A prospective randomized controlled trial was performed in SJL mice exposed to the myelin-related oligopeptide PLP_139-151_; a relapsing-remitting Experimental Autoimmune Encephalitis (EAE) model. 100 mice were randomized into five groups of 20. Group 1-Unimmunized prior to disease induction. Group 2-Immunized twice with a 74 kDa fusion protein vaccine against MAP; delivered 28 days and 7 days prior to disease induction. Group 3- 1 dose of 74 kDa vaccine 10 days post-disease induction. Group 4- Attenuated whole cell MAP vaccine (ΔSigH) 28 days prior to disease induction. Group 5- ΔSigH vaccine 10 days post disease induction.

Disability was quantified using a EAE disability scoring reference ranging from 0 to 5.

Significant decreases in peak disability were seen in the bimodal peaks of this relapsing-remitting model 38% (p<0.006) and 40% (p<0.001). ΔsigH immunized mice lost half as much weight as controls post disease induction. The results suggest that environmental MAP antigen exposure may play an etiologic role in the development of EAE.

## Introduction

The sine qua non of Multiple Sclerosis (MS) involves immune-mediated destruction of myelin in the central nervous system resulting in scarring, abnormal conductivity, and resulting disability. There are two forms of the disease. Relapsing-remitting MS accounts for 80% of the disease burden whereas the primary progressive form accounts for 20%. Twin studies show concordance for MS between identical twins to be 30% and 3% between fraternal twins [1]. This suggests contributions from genetics as well as the environment.

Genetic evaluations have shown increased risk of MS with haplotypes HLA-A as well as HLA-DRB1, the combination of which leads to a 23 fold increase [2]. Known predisposing factors, which may work through epigenetic mechanisms, include low vitamin D levels and aberrant response to Epstein Barr Virus (EBV) infection. Vitamin D deficiency has been associated with a two fold increased risk of MS [3]. High levels of Epstein Barr Nuclear Antibody (EBNA) confer a thirty six fold increase in risk [4].

Kurtzke et.al. detailed an epidemiologic case for infection as that environmental agent [5]. Investigation of MS clusters have revealed some clues. One, in the Faroe Islands, highlighted the rise in MS incidence years after British military occupation [5] and another, in Galion Ohio, revealed an increased MS incidence after a soil giveaway associated with a new high school gymnasium construction [5]. These suggest a possible fastidious organism, which has been further bolstered by studies associating Mycobacterium avium *ssp. paratuberculosis* with MS. Cossu et. al. reported higher humoral titers to MAP in M.S. patients [6] and Mameli et. al. reported increased serum and CSF reactivity to MAP antigens in patients as well [7]. With respect to EAE murine models, an increased genetic susceptibility to mycobacterial species has been shown [8] and mycobacterial cage contamination can be present [9].

Prior Epstein Barr Virus (EBV) has been found to be present in 100% of MS patients [10]. The high level of seroconversion of non-MS patients suggests that it is likely necessary, but not sufficient to cause Multiple Sclerosis. EBV may lay the groundwork for a second hit to occur resulting in MS. Establishing causation for a second-hit agent may be problematic if we adhere to Koch’s Postulates. If, for example, the second hit induces a molecular mimicry with myelin antigens then immunologic clearance of the agent may occur leaving little trace. An alternate way of establishing causation would be to immunize against the infectious agent and see if this prevents the disease. Ristori et.al. have reported some success in treating MS with BCG [11], which may act as a crude vaccine for MAP, just as Cowpox was a crude but effective vaccine against Smallpox in the late 1700’s.

We have set out to immunize against MAP specifically in EAE mice to see if this significantly decreases neurologic disability. Our findings suggest a large and significant improvement in mice which were immunized against MAP prior to disease induction.

## Methods

### ΔsigH vaccine preparation

A live attenuated M. avium ssp. *paratuberculosis* vaccine was prepared by Dr. Talaat’s lab as described in Shippy et. al. [12]. The sigH regulator gene was altered to decrease MAP pathogenicity.

### 74 kDa. fusion protein preparation

The 74 kDa. fusion protein vaccine was constructed de novo at Proteos Labs (Kalamazoo, MI) from the DNA sequence provided by Dr. Chang’s lab in Chen et. al. [13]

### SJL mice immunization

**Day -28** Cryo-vials on dry ice from Proteos containing 0.5 mg/ml Map74F in TBS, pH 7.4 received. The vaccine was stored at −80°C.

**Day -22** 100 SJL mice (Envigo, male, PO #193687, R #3778, 5-6 weeks old) were received, individually examined and housed in ten filter-topped cages of ten mice each in a HEPA-filtered enclosure. The mice were placed in quarantine with daily inspections.

**Day -21** The mice were ear tagged (SOP 810) with unique numbers for identification purposes, and weighed (RESULTS) and sorted into five treatment groups of 20 mice each based upon average body weight.

**Day -15** Received via Federal Express on coolants from the University of Wisconsin in 50 ml conical tubes and stored at 4-8°C

**Day 0** Group 1: 2 ml sterile PBS (Sigma, Cat. D8537, lot RNBR3321) was added to one vial Ribi adjuvant system (RAS, Sigma, Cat. S6322, lot 125M4172V) which had been warmed to 40°C. The mixture was vortexed for 3 minutes, inverted and vortexed for an addition minute.

Group 2: RAS was warmed to 40°C prior to addition of 2 ml (2 vials) Map74F (equilibrated to room temperature). The mixture was vortexed for 3 minutes, inverted and vortexed for an addition minute.

Group 4: 1 ml of the SigH was diluted into 1 ml sterile PBS.

The mice were weighed (RESULTS). The mice in Groups 1,2,4 were injected subcutaneously (SC, SOP 1610) with 0.1 ml of their respective vaccine.

**Day 21** Group 1: 2 ml sterile PBS was added to one vial RAS which had been warmed to 40°C. The mixture was vortexed for 3 minutes, inverted and vortexed for an addition minute.

Group 2: RAS was warmed to 40°C prior to addition of 2 ml (2 vials) Map74F (equilibrated to room temperature). The mixture was vortexed for 3 minutes, inverted and vortexed for an addition minute. The mice were weighed (RESULTS). The mice in Groups 1, and 2 were injected SC with 0.1 ml of their respective vaccine.

**Day 28** 15 mg PLP_139-151_ (BioSynthesis, lot T6928) was dissolved in 15 ml sterile PBS to prepare a 1 mg/ml solution. 100 mg heat-killed *Mycobacterium tuberculosis* H37 RA (Difco, Cat. 231141) was homogenized in 50 ml squalene (Sigma, Cat. S3626, lot 118K0847) to prepare a 2 mg/ml suspension (Complete Freund’s Adjuvant, CFA) 50 μg pertussis toxin (Sigma, Cat. P7206, lot SLBR6905V) was dissolved in 50 ml sterile PBS to prepare a 1 μg/ml solution. 11 × [1ml PLP was homogenized with 1 ml CFA] on an ice bath. A droplet of the emulsion floated on water did not disperse. The mice were weighed and injected SC in both flanks with 0.1 ml PLP/CFA emulsion. One hour later the mice were injected by intraperitoneal route (IP, SOP 1630) with 0.2 ml pertussis toxin (200 ng/mouse).

**Day 30** 50 μg pertussis toxin was dissolved in 50 ml sterile PBS to prepare a 1 μg/ml solution.

The mice were injected IP with 0.2 ml pertussis toxin (200 ng/mouse).

**Day 38** RAS was warmed to 40°C prior to addition of 2 ml (2 vials) Map74F (equilibrated to room temperature). The mixture was vortexed for 3 minutes, inverted and vortexed for an addition minute.

Group 5: 1 ml of the SigH was diluted into 1 ml sterile PBS.

The mice were weighed (RESULTS). The mice in Groups 3 and 5 were injected subcutaneously (SC, SOP 1610) with 0.1 ml of their respective vaccine.

**Days 29-67** The mice were weighed (RESULTS) and scored daily for signs of disease as in TABLE 1.

**Table 1.**
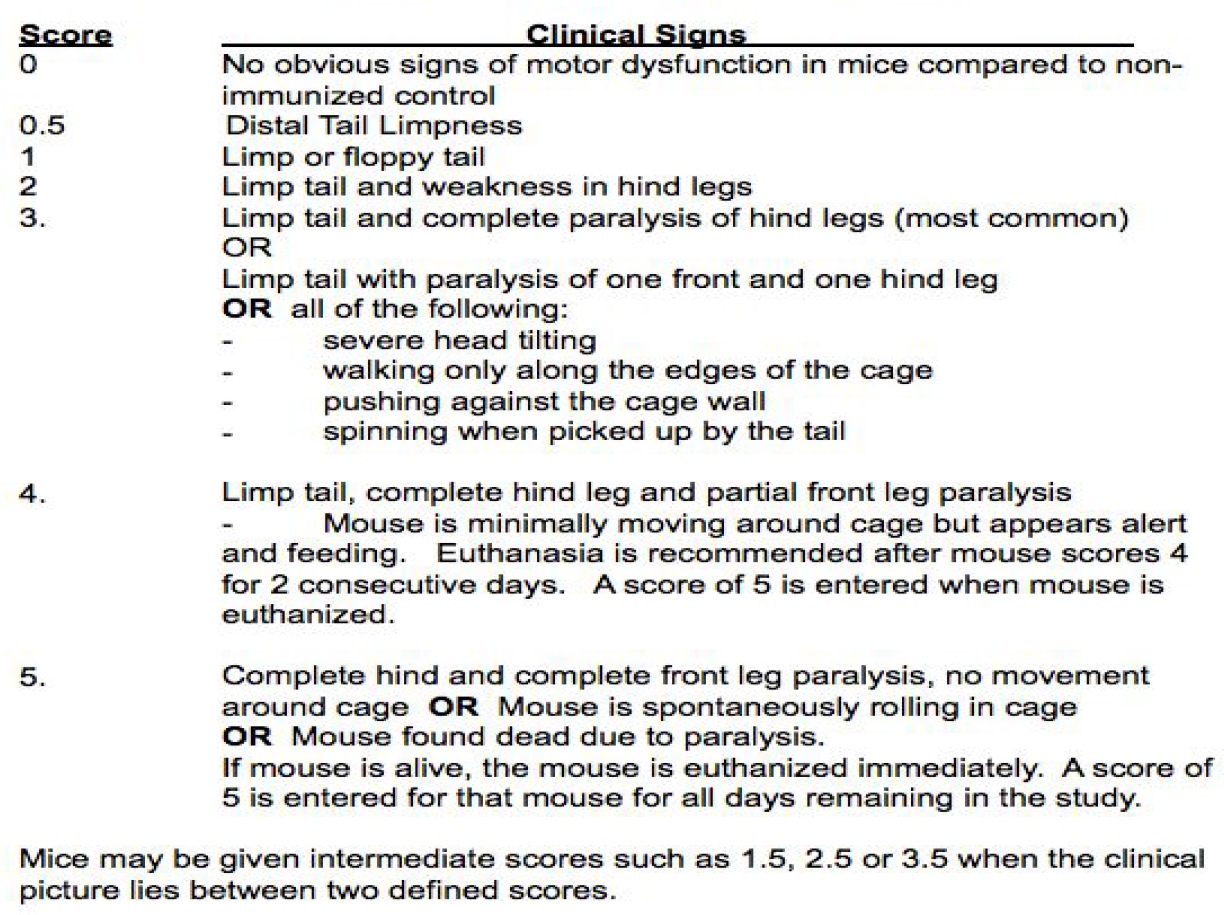
EAE SCORING PARAMETERS.

### Statistical Analysis

Daily disability scores were documented and significance (p-value) was calculated vs Group 1 by two-way ANOVA followed by Bonferroni post test analysis.

## Results

Vaccination targeted against MAP was effective in providing significant relief from neurologic disability in the EAE mouse model of Relapsing-Remitting Multiple Sclerosis. Peak disability, as measured by the EAE Scoring Parameters, was 38% less (p<0.006) in the SigH immunized mice for the first flare of disability and 40% less (p<0.001) for the relapse (Fig. 1).

**Fig. 1.**
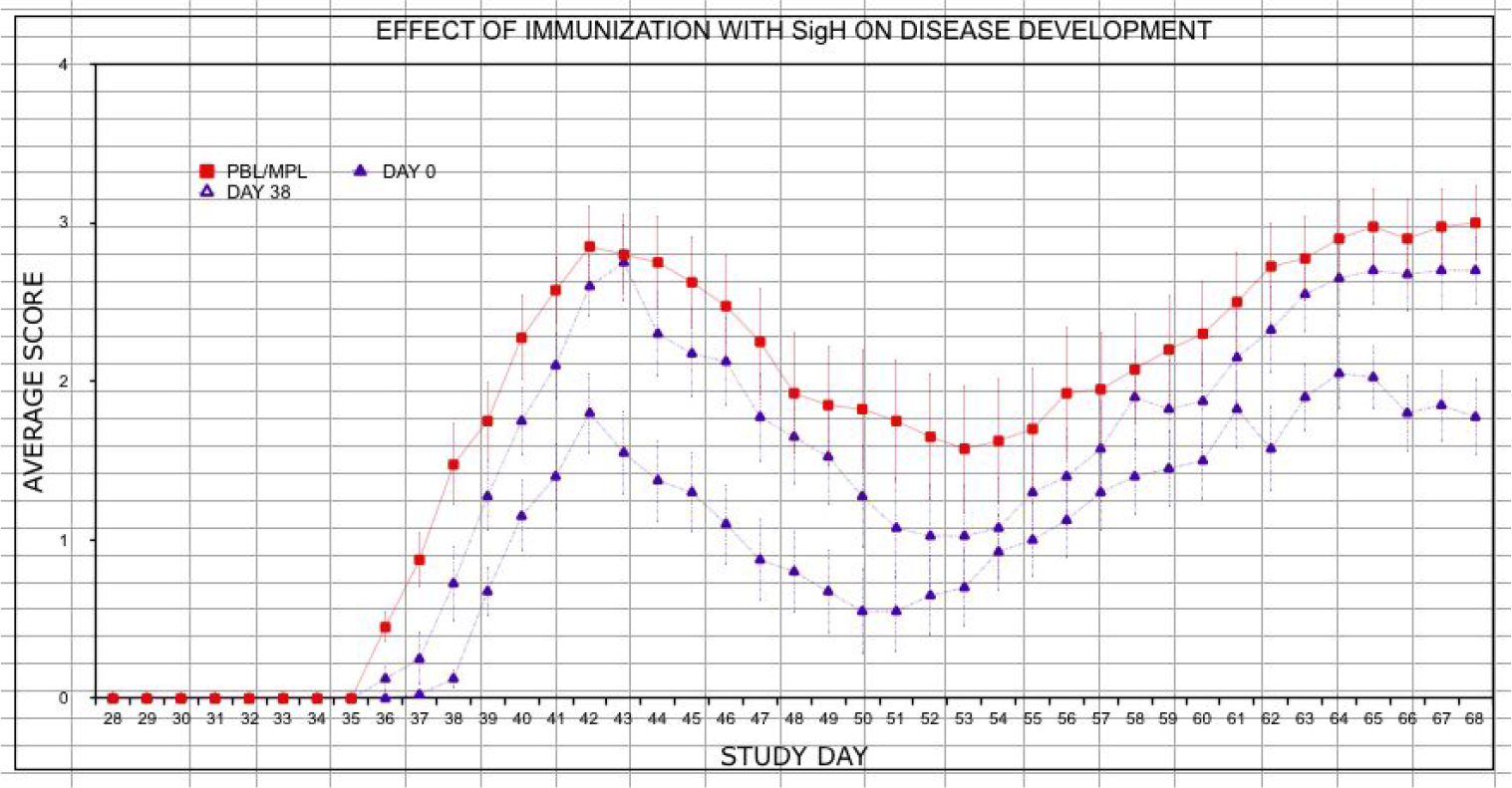

The 74 kDa Fusion vaccine resulted in a 24% reduction (p<0.03) in the first flare of disability and a 23% (p<0.05) for the relapse (Fig. 2). Late vaccination with either vaccine, once clinical disease onset occurred, did not result in statistically significant decreases in disability.

**Fig. 2.**
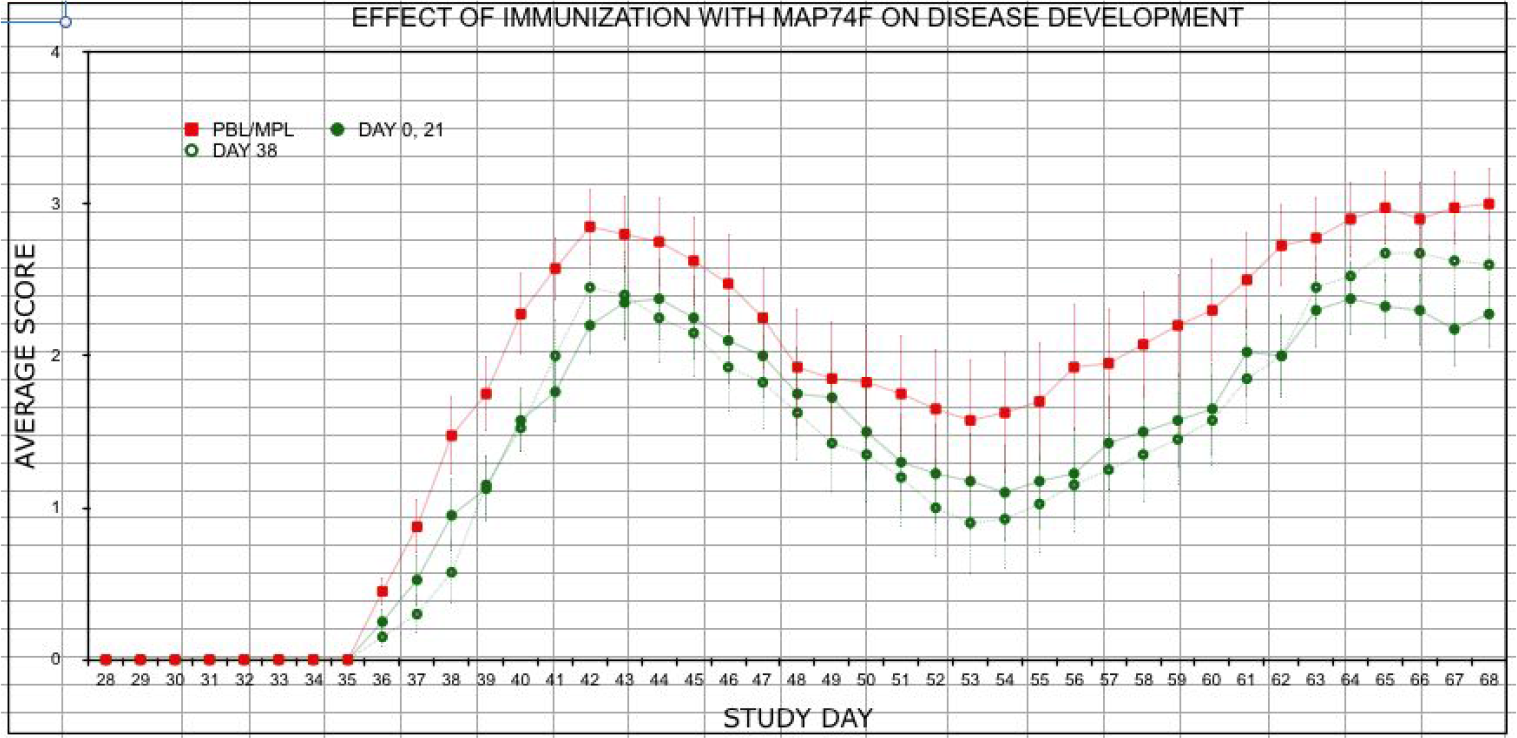

In addition to improvement in disability scores, mice immunized with the ΔsigH vaccine, either before or after disease induction, exhibited significantly less weight loss (p<0.05, p<0.05) compared to pre-disease induction weights. −11.56% for Control Group 1, −4.5% for Group 4, and −5% for group 5. (Fig. 3)

**Fig. 3.**
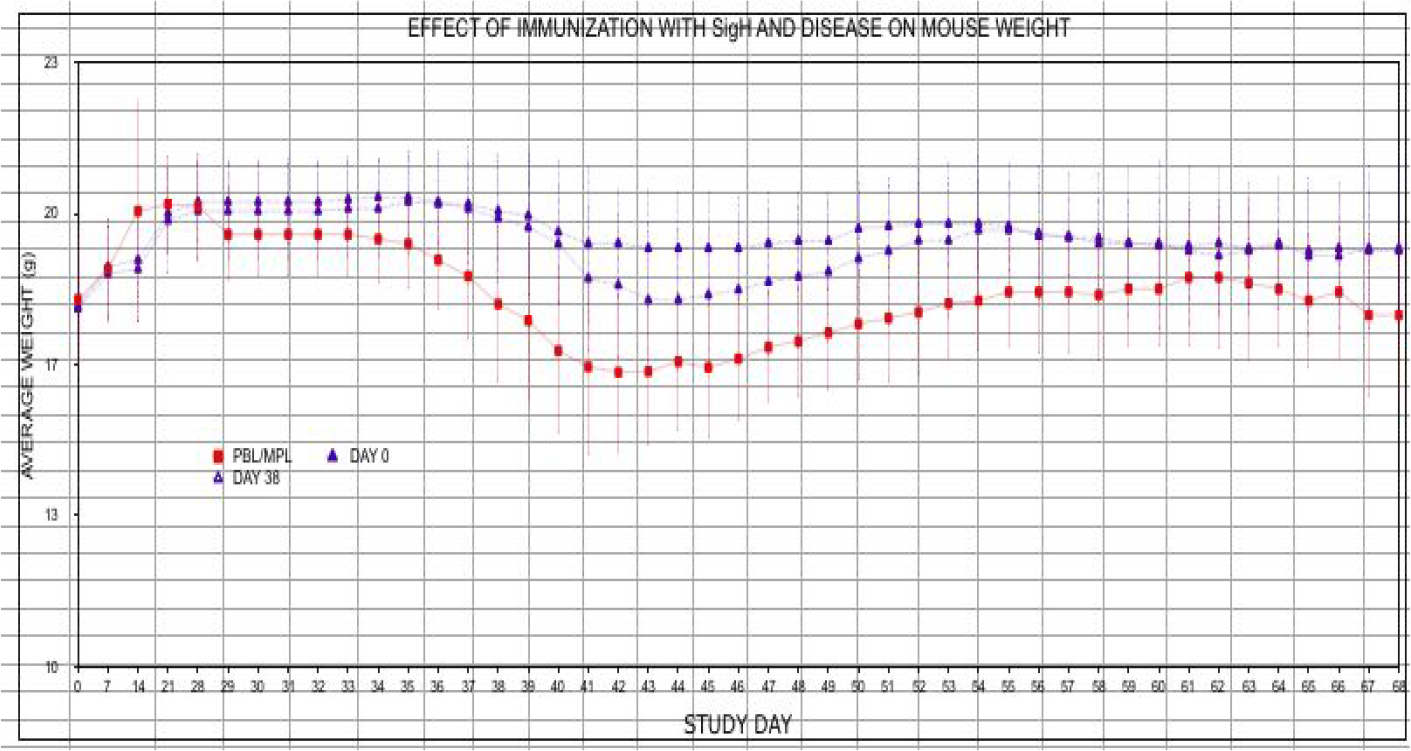

## Discussion

These findings represent an evolution from examining associations between multiple sclerosis and MAP to establishing a causative role for such agents. Establishing causation results in a myriad of actionable interventions to assist those suffering from MS.

MAP’s presence can be found in the dairy industry as a cause for Johne’s disease, an autoimmune inflammatory intestinal condition. Widespread vaccination does not occur to date. Pasteurized milk can contain viable MAP as well as its epitopes [14]. Subsequent contamination of groundwater and soil occur with MAP, and it is represented in biofilms in municipal water supplies [15]. An effective bovine vaccination program and further research on destroying biofilms could modify the risk of developing MS or its relapses.

Focused human vaccination for susceptible individuals, those with a HLA-DRB1 haplotype and/or those with elevated EBV antibodies, would also be an option. Support is further suggested by the Ristori et. al. finding that *BCG* can decrease MS lesions on MRI. [11] Formerly thought to be a nonspecific immune system booster, *BCG* and it’s mycobacterial antigens may represent partial vaccination against MAP. If EBV is considered the first hit and MAP is considered the second hit in this pathogenic mechanism, it follows that EBV vaccination efforts continue in earnest.

Apart from environmental containment efforts and vaccination, a third avenue of treatment might focus on inducing tolerance to MAP antigens at a young age. An environmental cline exists in MS epidemiology which reveals a higher risk of developing MS as distance from the equator increases.(Fig.4)

**Fig. 4.**
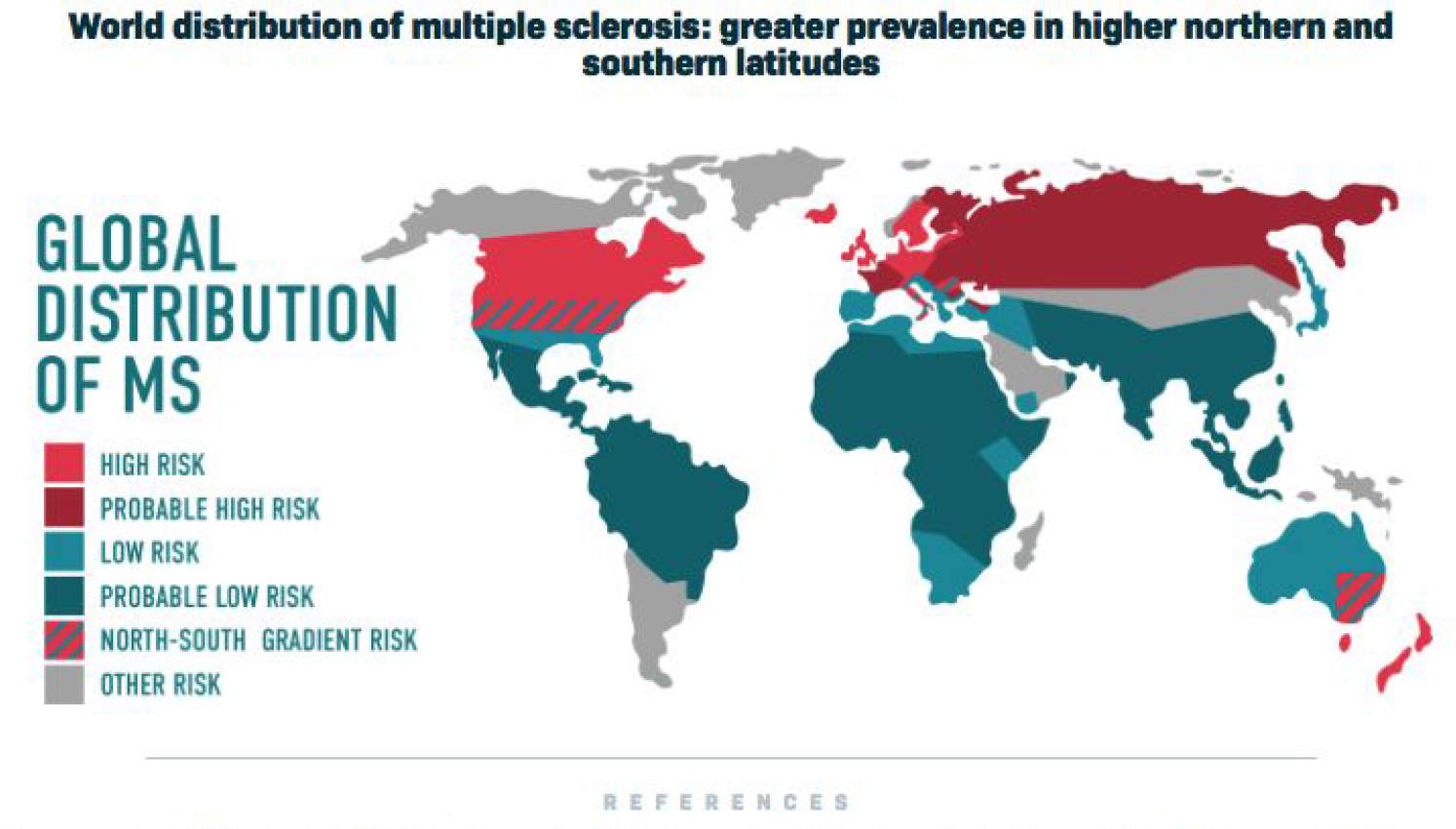

It has also been shown that moving from a low-incidence area to a high-incidence area after age fifteen will result in an individual having a MS risk equivalent to that of the low-risk prior address [16,17] Whereas, an individual moving from a high to low incidence area will retain that higher risk if that person is older than fifteen. This may act as an epidemiologic clue linking EBV and MAP as the average age of seroconversion to EBV is the teen years [18]. It follows that exposure and subsequent tolerance to M. Avium prior to EBV infection may be protective whereas EBV acquisition in a susceptible individual prior to MAP exposure may be pathologic. Support for an exposure cline to match the MS disease cline can be traced to US military recruit data from the 1950’s [19]. A PPD study using M.avium proteins revealed higher rates of positivity in Southern recruits (70%) relative to their Northern counterparts (20%).

Overall, these findings highlight the need to systematically investigate suspected infectious triggers for MS and other autoimmune diseases with the goal of not merely uncovering associations but making an attempt to establish causative links through vaccination. Establishing these links would open the door to antiinfectives, vaccination, and perhaps immunotherapy. All of which would get closer to prevention than the current medication options, which reflect downstream damage control and are prohibitively expensive.

## Acknowledgements

Funding for this research made possible by a grant from Jim and Shirley Balk/Spectrum Health Foundation.

